# Triggering closure of a sialic acid TRAP transporter substrate binding protein through binding of natural or artificial substrates

**DOI:** 10.1101/2020.12.02.404004

**Authors:** Martin F. Peter, Christian Gebhardt, Janin Glaenzer, Niels Schneberger, Marijn de Boer, Gavin H. Thomas, Thorben Cordes, Gregor Hagelueken

## Abstract

The pathogens *Vibrio cholerae* and *Haemophilus influenzae* use tripartite ATP-independent periplasmic transporters (TRAPs) to scavenge sialic acid from host tissues. They use it as a nutrient or to evade the innate immune system by sialylating surface lipopolysaccharides. An essential component of TRAP transporters is a periplasmic substrate binding protein (SBP). Without substrate, the SBP has been proposed to rest in an open-state, which is not recognised by the transporter. Substrate binding induces a conformational change of the SBP and it is thought that this closed state is recognised by the transporter, triggering substrate translocation. Here we use real time single molecule FRET experiments and crystallography to investigate the open- to closed-state transition of VcSiaP, the SBP of the sialic acid TRAP transporter from *V. cholerae*. We show that the conformational switching of VcSiaP is strictly substrate induced, confirming an important aspect of the proposed transport mechanism. Two new crystal structures of VcSiaP provide insights into the closing mechanism. While the first structure contains the natural ligand, sialic acid, the second structure contains an artificial peptide in the sialic acid binding site. Together, the two structures suggest that the ligand itself stabilises the closed state and that SBP closure is triggered by physically bridging the gap between the two lobes of the SBP. Finally, we demonstrate that the affinity for the artificial peptide substrate can be substantially increased by varying its amino acid sequence and by this, serve as a starting point for the development of peptide-based inhibitors of TRAP transporters.

## Introduction

The tripartite-ATP independent-periplasmic (TRAP) transporters are a large class of membrane transporters that are widespread in bacteria and archaea. They have unique properties in being a structural and functional mix of ATP-binding cassette (ABC) transporters and secondary-active transporters, both of which are well studied. TRAPs translocate a large variety of small substrates, most of which contain a carboxylic acid group ^1–3^. They rely on a periplasmic high-affinity substrate binding protein (SBP, in TRAP transporters also termed P-domain) that is assumed to scavenge and deliver substrate molecules to the actual transporter domains in the inner membrane ^4, 5^. For TRAP transporters, the latter are known as Q- and M-domains and consist of 4 and 12 transmembrane (TM) helices, respectively. In some representatives, the Q- and M-domains are fused into one peptide chain by an additional TM-helix ^5^.

The best studied TRAP transporters are currently the *N*-acetylneuraminic acid (commonly known as sialic acid or Neu5Ac) transporters HiSiaPQM and VcSiaPQM from the two human pathogens *Haemophilus influenzae* and *Vibrio cholerae* ^4, 5^. For these two pathogens, the TRAP transporter is the only uptake route for the exogenously synthesized sialic acid, which is present in host tissues. After uptake, the small sugar molecule is used to sialylate the bacterial lipopolysaccharides as a strategy to evade detection by the hosts innate immune system ^6–8^. To date, the structures of the transmembrane domains are unknown, but, high-resolution crystal structures of the open form of the P-domains of HiSiaP and VcSiaP and of the closed form of HiSiaP have been determined ^9–11^.

Like other SBPs ^12^, the TRAP transporter P-domains have a characteristic bilobal fold with a large substrate binding cleft. The two lobes are connected by a long α-helix, which forms the backbone of the structure. Substrate binding induces large conformational changes of the whole protein by more than 15 Å and the formation of a characteristic kink in the backbone helix ^9^. The closing motion and the binding mechanism of the SBP have often been compared to a Venus flytrap ^12, 13^. A PELDOR spectroscopy study (pulsed electron double resonance, also known as DEER) on spin labelled VcSiaP indicated that in frozen solution and without substrate, the P-domain “lurks” in the open state, similar to some ABC transporter SBPs ^14–20^. The substrate binding cleft has two highly conserved arginine residues (Arg125 and Arg145 in VcSiaP), which coordinate the carboxylic acid group of the substrate ^10, 11, 21^. In addition, an intricate water network around the bound substrate is an important determinant of substrate affinity ^22^.

Still, numerous questions concerning the role of the P-domains for TRAP transporters and concerning SBPs in general remain unanswered. Most importantly, it is still unclear how the sialic acid triggers the transition to the closed state. Considering that TRAP SBPs are potential drug targets, it is of high interest to find the requirements and specifics of a substrate that can initiate the transition.

Here, we use a combination of single-molecule FRET spectroscopy and X-ray crystallography to study the conformational changes of single VcSiaP molecules in real time and to investigate the molecular details of the closing mechanism.

## Results

### Open-closed dynamics of VcSiaP at room temperature

Binding of the substrate to the periplasmic SBP is the first step of the proposed mechanism for TRAP transporters and involves large scale conformational changes of the protein ^5^. PELDOR/DEER experiments on frozen VcSiaP solutions indicated that the protein switches its conformation in a substrate dependent manner ^18^. Here, we used single-molecule Förster-resonance energy transfer (smFRET) spectroscopy to investigate this important step of the mechanism by real-time measurements of VcSiaP dynamics under more physiological conditions. Such experiments should allow comparisons to SBPs from the ABC transporter family ^14, 17^. For this purpose, we employed the Q54C/L173C double mutant of VcSiaP that was used for the PELDOR/DEER experiments ^18^. The mutant protein was labelled with maleimide-derivatives of AlexaFluor 555 (donor) and AlexaFluor 647 (acceptor). The distance between both label positions was expected to decrease upon ligand binding, i.e., the transition from the open-to closed state should lead to a change from intermediate to high FRET-efficiency. On average we obtained a labelling efficiency > 90% with ~1.83 fluorophores per protein (0.68 donors and 1.15 acceptors) for Alexa555/647 and a resulting yield of donor-acceptor containing VcSiaP of ~49% (with 10% donor only and ~41% acceptor only molecules).

With this at hand, we studied the substrate dependence of the conformational changes in VcSiaP (Q54C/L173C) using solution-based FRET. This allowed us to simultaneously check whether the addition of the fluorescence label had any impact on the affinity to Neu5Ac and which conformational states of VcSiaP are seen in solution at room temperature. The relative populations of the open- and closed-conformation were determined in a titration experiment for different substrate concentrations, allowing us to determine a K_D_ value of 300 ± 100 nM (Fig 1A-C). This value was in very good agreement with a previously published ITC experiment ^11^ and a control ITC experiment for our particular wild-type construct (Figure 1E). Thus, the presence of the fluorescence labels does not influence the Neu5Ac affinity and the conformational changes in VcSiaP are directly coupled to the substrate binding to the protein (Figure 1DE).

**Figure 1:**
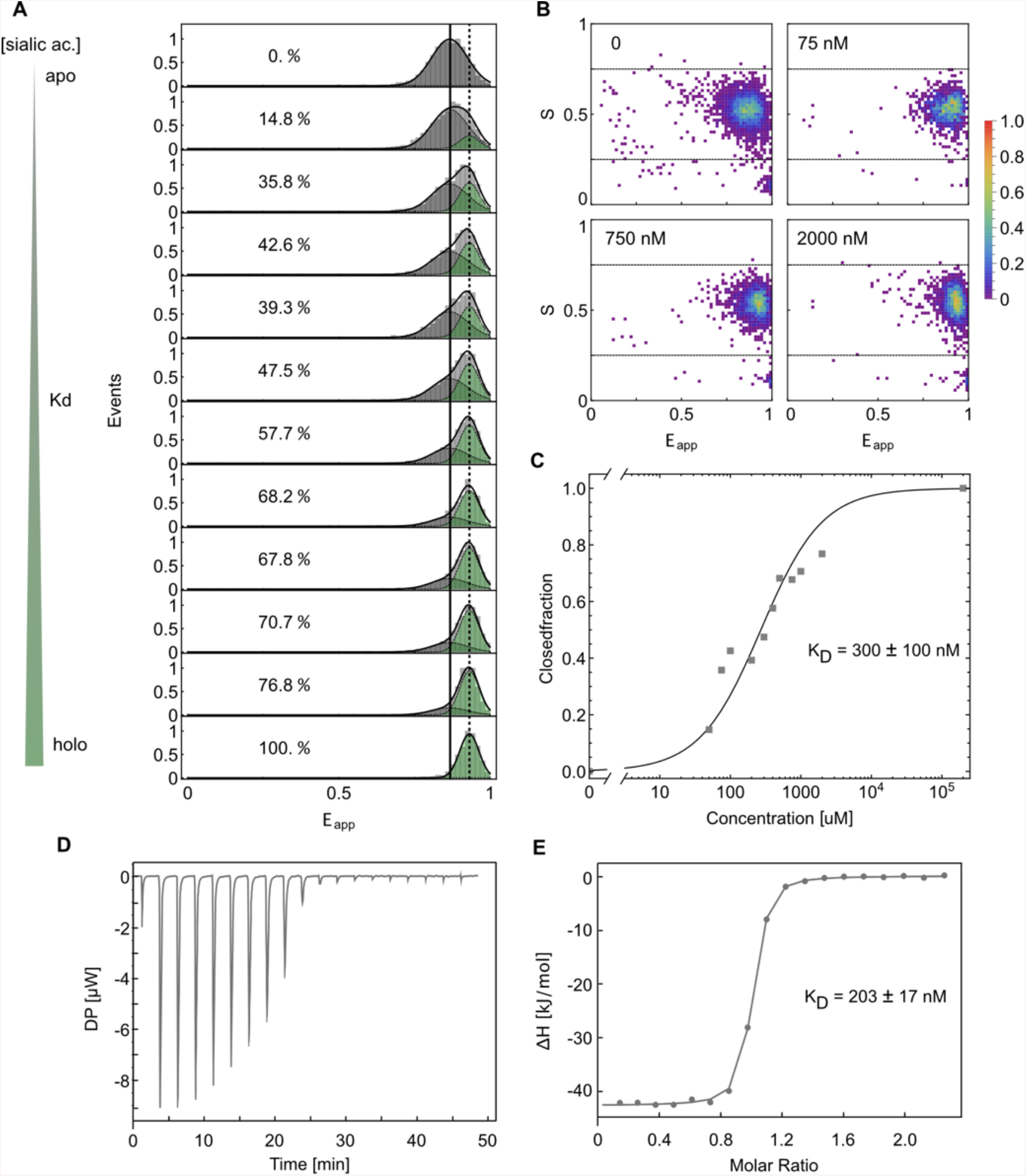
Steady-state Neu5Ac binding by VcSiaP. **A)** Apparent FRET efficiency histograms of VcSiaP with labels Alexa555/647 for different Neu5Ac substrate concentrations (0 (top) to 200 μM (bottom)). The fraction of molecules in the closed state *r*_*c*_ was determined by a global fit of 2 Gaussians for the open and closed state, shown in dark gray and green, respectively. **B)** Representative 2D ES-histograms from data in A) at 4 different concentrations. **C)** Fraction of closed VcSiaP *r*_*c*_ depending on ligand concentration extracted from Gaussian fits to data in A). The data points were fitted to the function 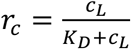 with a dissociation constant of K_D_= 300 *±* 100 nM (95% confidence interval). **D+E)** ITC data of Neu5Ac titrated to VcSiaP solution (left) and corresponding binding isotherm (right).

To derive the binding mechanism of VcSiaP, we performed smFRET experiments with surface-immobilized protein via scanning confocal microscopy. For this purpose, we attached VcSiaP to a glass-surface using its N-terminal His_6_-tag (Figure 2A). We analysed the conformational dynamics between open- and closed-state in real time using apo conditions (no ligand), 500 nM Neu5Ac (~K_D_) and 1 mM Neu5Ac (saturated ligand concentration). The FRET-states observed of VcSiaP (Q54C/L173C) at apo and saturated ligand conditions are fully compatible with diffusion-based experiments (Figure 3) and show the population of the open (absence of substrate) and closed (1 mM Neu5Ac) state (Figure 2BC). For concentrations close to the K_D_-value we observed discrete and stochastic transitions between low- and high-FRET efficiency states (Figure 2B, middle panel). The low and high FRET states have an average apparent FRET efficiency of 0.791 ± 0.012 (95% confidence interval) and 0.895 ± 0.010, respectively. They are similar to the average efficiencies in the absence (0.807 ± 0.003) and presence of saturating (0.898 ± 0.003) concentrations of Neu5Ac (Figure 2BC). Therefore, the transitions can be interpreted directly as transitions between the open (low E*) and closed (high E*) conformation.

**Figure 2.**
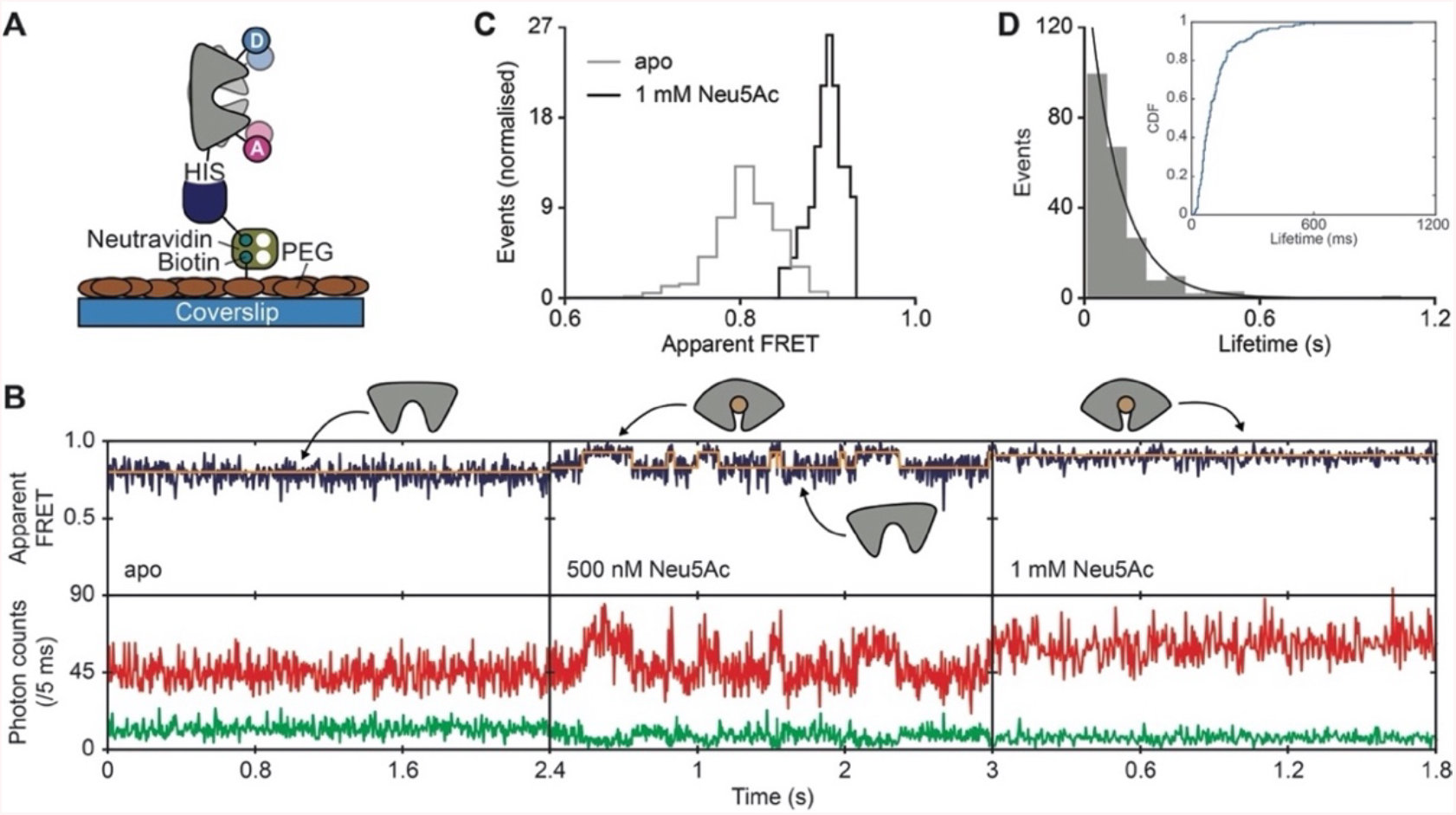
Study of immobilized VcSiaP proteins for analysis of sub-second dynamics. **A)** Schematic of surface immobilization procedure. **B)** Representative fluorescence trajectories of VcSiaP (Q54C/L173C) labelled with AlexaFluor 555 and AlexaFluor 647 under different conditions as indicated in the figure. The top panel shows the apparent FRET efficiency (blue) and donor (green) and acceptor (red) photon counts in the bottom panel. Orange lines are the most probable state-trajectory of the Hidden Markov Model (HMM). **C)** Apparent FRET efficiency histogram of all fluorescence trajectories recorded in the absence and presence of 1 mM Neu5Ac. **D)** Histogram and cumulative distribution function (CDF; inline panel) for the lifetime of the closed conformation (high FRET state) from all fluorescence trajectories recorded in the presence 500 nM Neu5Ac (N_traces_ = 45). The black line is an exponential fit.

**Figure 3:**
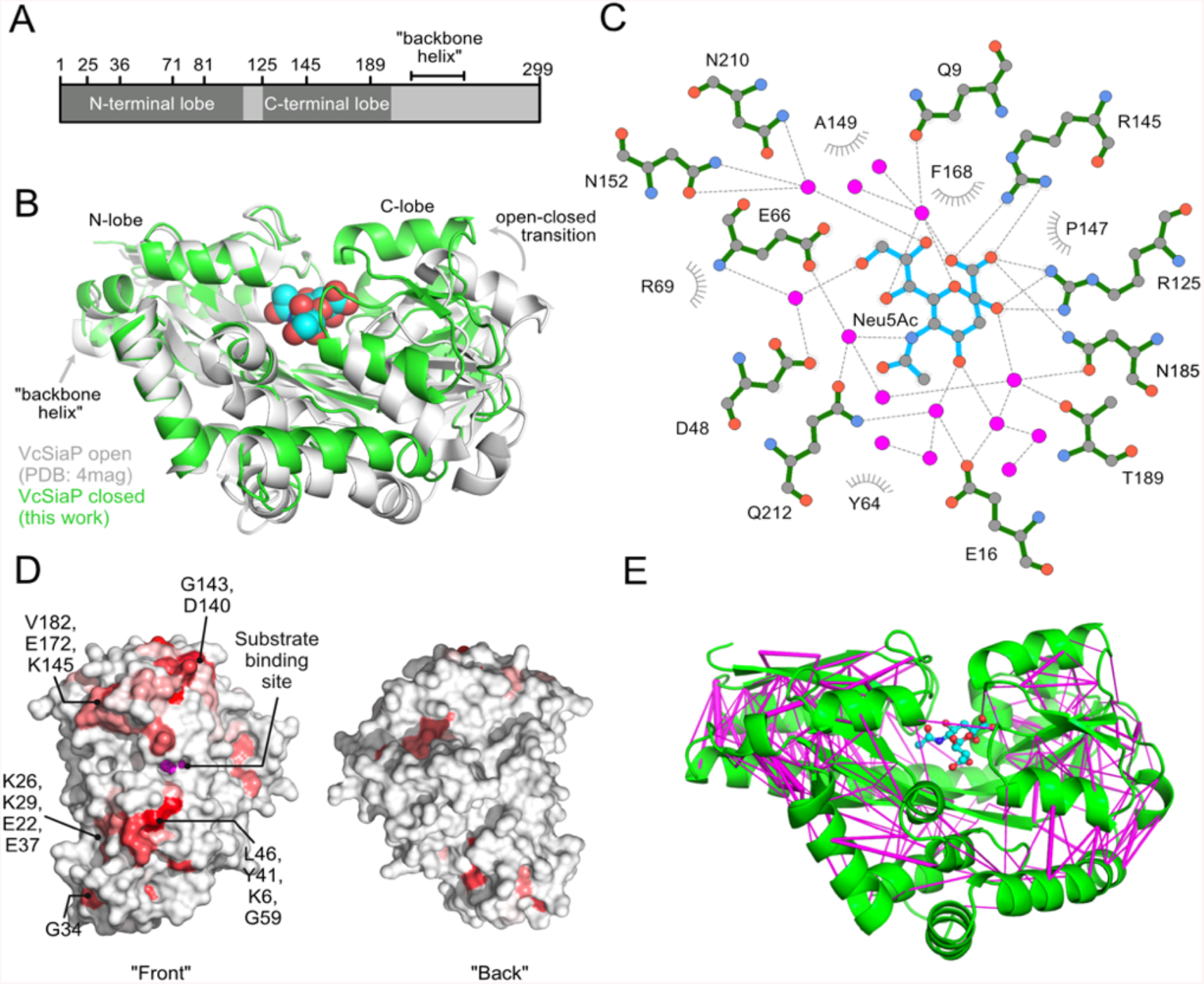
Structural classification of sialic acid-bound VcSiaP. **A)** Outline of the primary sequence of VcSiaP with structural motifs and positions of selected amino acids. **B)** Alignment of open (white, PDB-ID: 4MAG ^11^) and closed (green) conformations (this work) of VcSiaP. The substrate from the closed structure is represented as blue-coloured spheres. **C)** Ligplot analysis of the substrate binding interactions from VcSiaP. The water molecules are represented as magenta spheres ^50^. **D)** Surface conservation of TRAP transporter SBPs, calculated with the ENDscript server ^26^ and mapped onto the closed VcSiaP structure. The proteins that are used for this analysis are listed in a supporting file. The protein is shown as grey surface model and red regions indicate conserved surface regions. The two views are rotated by 180°. **E)** Evolutionary couplings ^28^ analysis of the closed VcSiaP structure as model. The protein is shown as a green cartoon and the bound substrate as blue-red ball-sticks. Interactions between pairs of residues that are identified as conserved from the analysis are shown as magenta lines.

To characterize the kinetics of the conformational transitions, we modelled the underlying state-trajectory of the fluorescence trajectories by a Hidden Markov Model (HMM ^23^) with two states, open and closed (Figure 2B; middle panel). From the state assignment of the HMM the individual lifetimes of the closed state were extracted. The distribution of these lifetimes was fitted with an exponential distribution and has an average lifetime of 122 ± 16 ms (95% confidence interval) (Figure 2D). Thus, once VcSiaP is in the closed conformation, it takes on average around 120 ms to re-open and release the ligand. Within the detection limit of smFRET and a total duration of almost 20 min recording time (850 fluorescence trajectories, i.e., 700 apo and 150 holo state), we observed that VcSiaP is exclusively in the open state when the ligand is absent and only closes when the ligand is bound (Figure 2B). Such a behavior would be compatible with an induced-fit (Venus flytrap) mechanism of ligand binding assuming that fluctuations between both conformations are not faster than the 5 ms time resolution we could achieve in the presented experiments. If conformational sampling was indeed (much) faster than this, other binding mechanism are still possible, where open-to-closed state transitions are not only driven by ligand.

### Crystal structure of substrate-bound VcSiaP

To create an atomic model of the closed state of VcSiaP that was observed in the smFRET experiments, we co-crystallised the protein with Neu5Ac. Single crystals were observed after three months of incubation at 20 °C. The crystal structure was solved at 1.6 Å resolution by molecular replacement ^24^, using the closed state of HiSiaP as the search model (PDB-ID: 3B50 ^10^). The electron density map clearly revealed a Neu5Ac molecule in the binding pocket, tightly bound by residues R125, R145 and N185 (Figure 3BC). The structure contains residues 1-299 of the mature protein (i.e. after cleavage of the signal peptide, Figure 3A) and was refined to R/Rfree values of 0.221/0.257 (Table 1). While the main interactions between protein and substrate are similar to other TRAP SBPs, subtle differences in the water network around the substrate were detected (SI Figure 1). These differences might explain the comparably low affinity of VcSiaP towards Neu5Ac ^11^. Overall, the structure is similar to the HiSiaP/Neu5Ac complex with an r. m. s. d. value of 0.7 Е for 279 Cα atoms. Still, the fact that the HiSiaQM membrane transporter discriminates between VcSiaP and HiSiaP *in vitro* ^25^ implies that important structural differences exist. We hence used the ENDscript server ^26^ to identify conserved areas on the surface of the closed state structure. The result is shown in Figure 3D (see SI for the underlying alignment). Clearly, residues surrounding the closed substrate binding cleft are strongly conserved. Considering the notion that it is the closed state of the SBP which is most likely recognised by the QM domains ^5^, this conserved and continuous surface region is likely involved in the P-QM interaction, as proposed for the YiaMNO TRAP transporter ^27^.

**Table 1:**
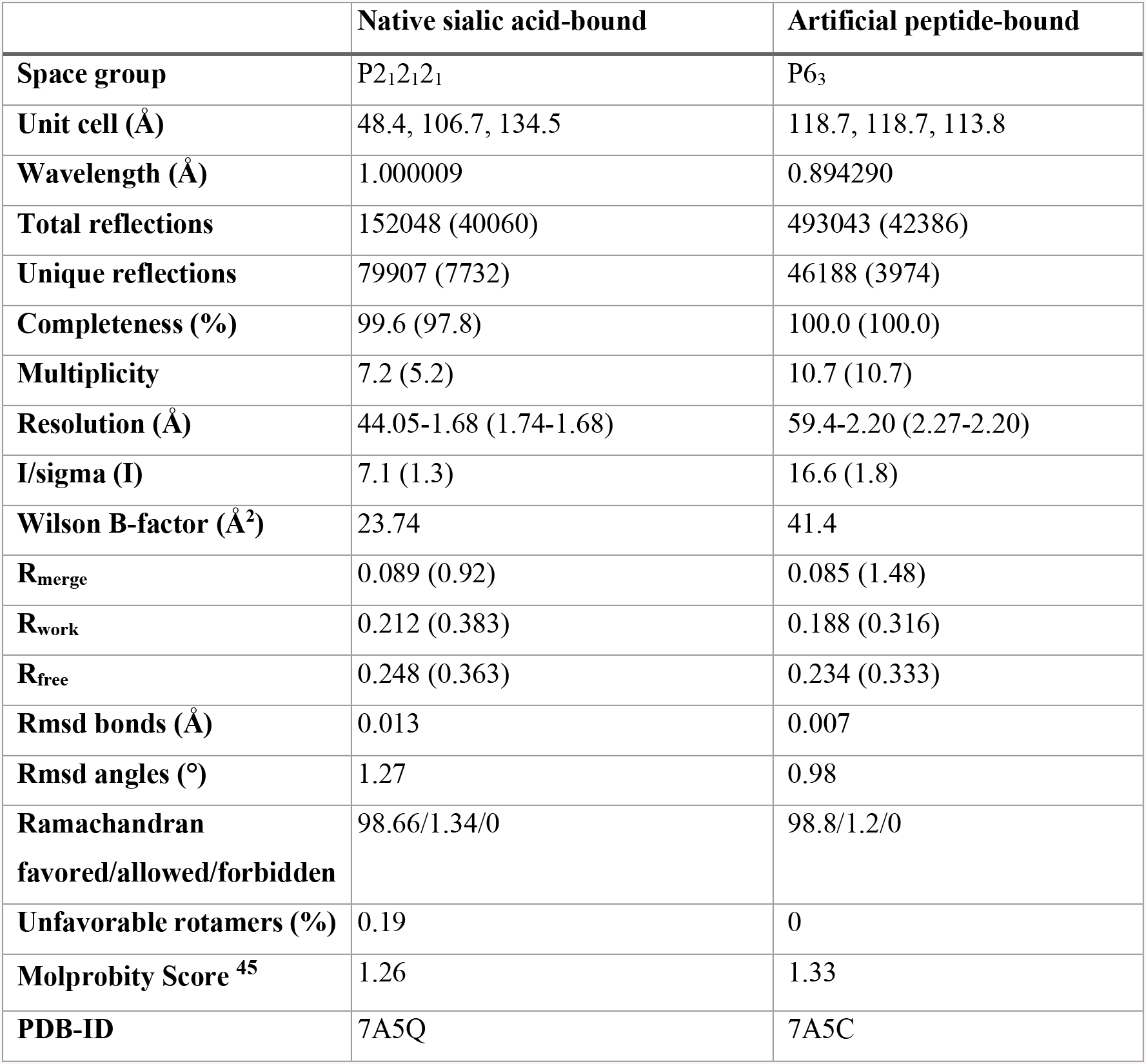
Data collection and refinement statistics.

Within this region, there was no obvious candidate that immediately explains the described discrimination between VcSiaP and HiSiaP, but, we noticed a surface patch with a number of reversed charges (D25, E28, K21, K36 in VcSiaP corresponding to K26, K29, E22, E37 in HiSiaP). This feature represents an interesting candidate for further experiments addressing this question (Figure 3D). The magenta lines in Figure 3E illustrate the spatial relation between conserved residue pairs of SBPs mapped onto the closed state structure of VcSiaP ^28^. Strikingly, although the two lobes clearly touch each other in the closed state (Figure 3D), there is only one weak evolutionary coupling of residues across the gap between the two lobes. Hence, it appears that the substrate is the “glue” that holds the two lobes of the SBP together, fitting nicely to the smFRET results above.

### A 9-mer peptide can artificially trigger SBP closing

From a structural and mechanistic perspective, it remains poorly understood how the large conformational change from the opened- to the closed state conformation is triggered. It has been proposed that a conserved triad of residues (R125, E184 and H207, VcSiaP numbering) in the hinge region is involved ^11^. But, our previous experiments have shown that these residues have no influence on the overall conformational change. ^18^ Mutants of these residues had a reduced affinity for Neu5Ac but were still able to adopt the closed-state structure. A serendipitous hint towards an answer to this question was obtained during our efforts to crystallize various substrate binding site mutants of VcSiaP with attached EPR nitroxide spin labels (R1). The crystal structure of one of them, a R125A mutant, was solved at a resolution of 2.1 Å by molecular replacement using the VcSiaP wild-type as search model (PDB-ID: 4MAG, ^11^) and refined to R/Rfree values of 0.199/0.232 (Table 1). Inspection of the initial electron density map surprisingly revealed a peptide in the substrate binding site (SI Figure 2). Also, the R125A mutation was clearly visible and the two R1 spin labels at positions 54 and 173 (far away from the peptide: SI Figure 3) were well defined in the electron density (SI Figure 2).

During refinement, it became clear that the peptide is connected to a neighboring molecule in the crystal and is in fact a part of the latter molecules’ affinity tag. Thus, two neighboring SBPs form an interlocked dimer in the hexagonal crystal packing (Figure 4A). The “tag peptide” is involved in many interactions with the SBP domain, especially at the entrance of the Neu5Ac binding cleft (Figure 4B). The N-terminal His_6_-tag itself is not visible in the electron density and therefore apparently disordered. Interestingly, position D-14 of the tag peptide interacts with R145 of the SBP. The latter is slightly disordered in our structure, but all known substrates of TRAP transporters interact with this residue ^21^. Further, there are several interactions between Y_−15_ of the tag peptide and R69 of the SBP, T-10 of the tag peptide and S42 of the SBP as well as multiple hydrophobic interactions (Figure 4B). As mentioned above, the crystal structure is of the R125A Q54R1 L173R1 mutant of VcSiaP. Modeling the arginine residue back into the structure shows that its presence would not interfere with the tag peptide binding site and DLS measurements show a similar behavior in dimerization between the tagged wildtype and R125A mutant (SI Figure 3). We used difference distance matrices to analyze the conformational change induced by peptide binding and compared them to the changes induced by Neu5Ac binding (Figure 4C). This comparison reveals clear signs of closing of the peptide-bound SBP. Although the scale of the two half-matrices is different, their overall pattern is strikingly similar. Thus, the open to Neu5Ac-bound movement and the open to peptide-bound movement are closely related with the same rigid bodies moving relative to each other, albeit on different length scales. It appears that the peptide, a completely artificial substrate, has triggered closing and its presence has trapped a previously undescribed intermediate state of the SBP.

**Figure 4:**
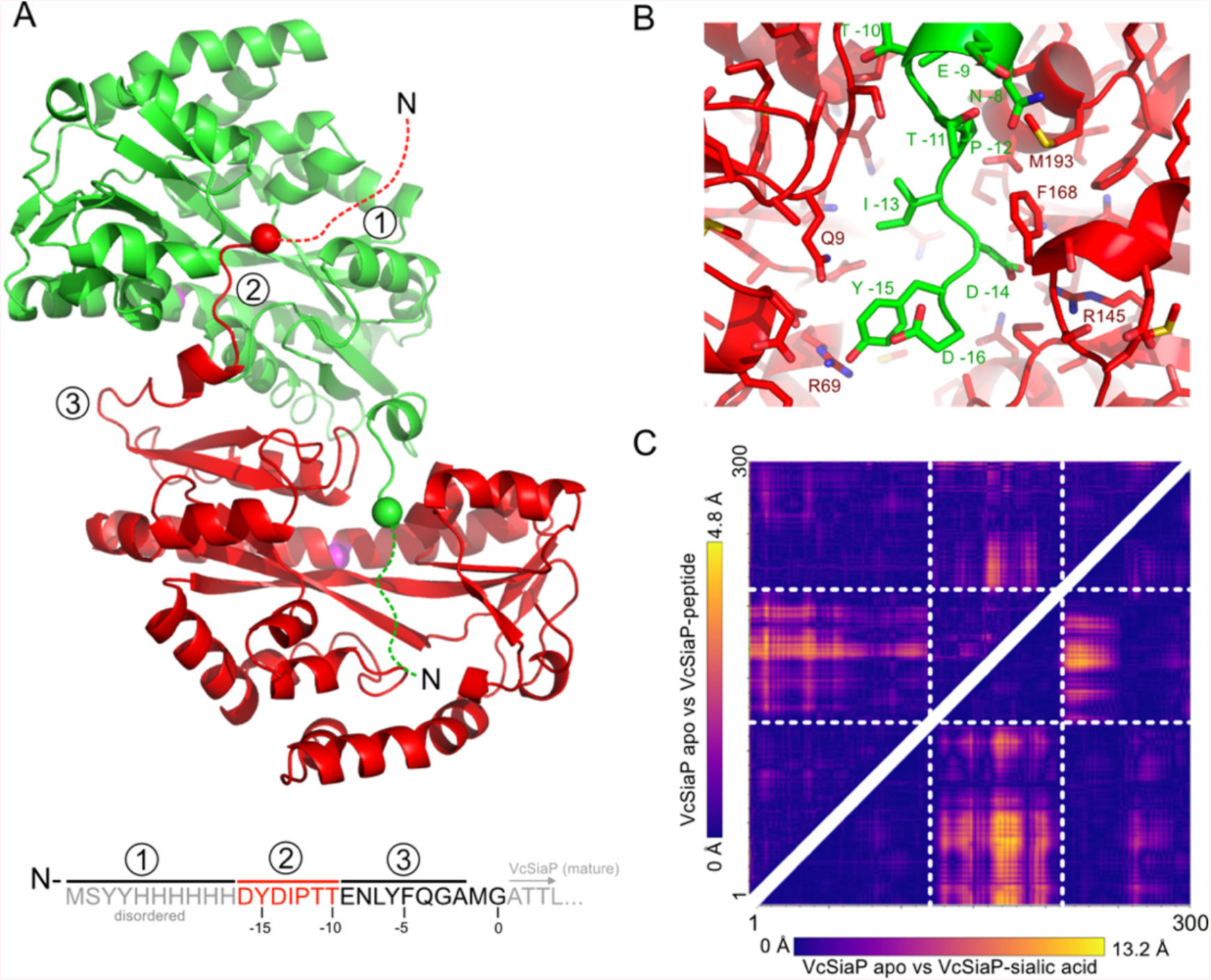
Crystal structure of an intertwined dimer of VcSiaP. **A)** Two interlocked monomers of VcSiaP are shown in cartoon representation (green and red). The N-terminus of one domain is bound to the substrate binding cleft of the other domain. The N-termini are indicated by spheres. Numbers in circles and the amino acid sequence at the bottom clarify, which part of the N-terminal tag is bound to the neighboring molecule. The magenta sphere indicates residue Q245, which was used to mutate and attach a fluorescence label for binding studies (see below). **B)** Close-up of the peptide-VcSiaP interactions in the dimeric crystal structure. **C)** Difference distance matrices between the substrate-free and peptide-bound VcSiaP (top-left half) and the substrate-free and - sialic acid bound VcSiaP structures (bottom-right half). The coloring of the two matrices (different scales) indicates the extend of the conformational changes between each pair of structures. Rigid bodies are indicating by dashed lines. The matrices were created with mtsslWizard ^51^ (http://www.mtsslsuite.isb.ukbonn.de).

### Characterization and strengthening of the peptide-SBP interaction

The discovery that a peptide can bind to the SBP opens an interesting road towards the design of a peptide-based inhibitor for TRAP transporter SBPs. As a first step into this direction, we designed a peptide-scan, i.e. a cellulose membrane with a set of 9-mer peptides attached via spot synthesis ^29^. Binding of the SBP to these short 9-mer peptides was detected by a fluorescence labelled SBP, where the label was attached to an artificially introduced cysteine on the “backside” of the protein (Q245C-DyLight800). The results of these experiments are shown in Figure 5A.

**Figure 5:**
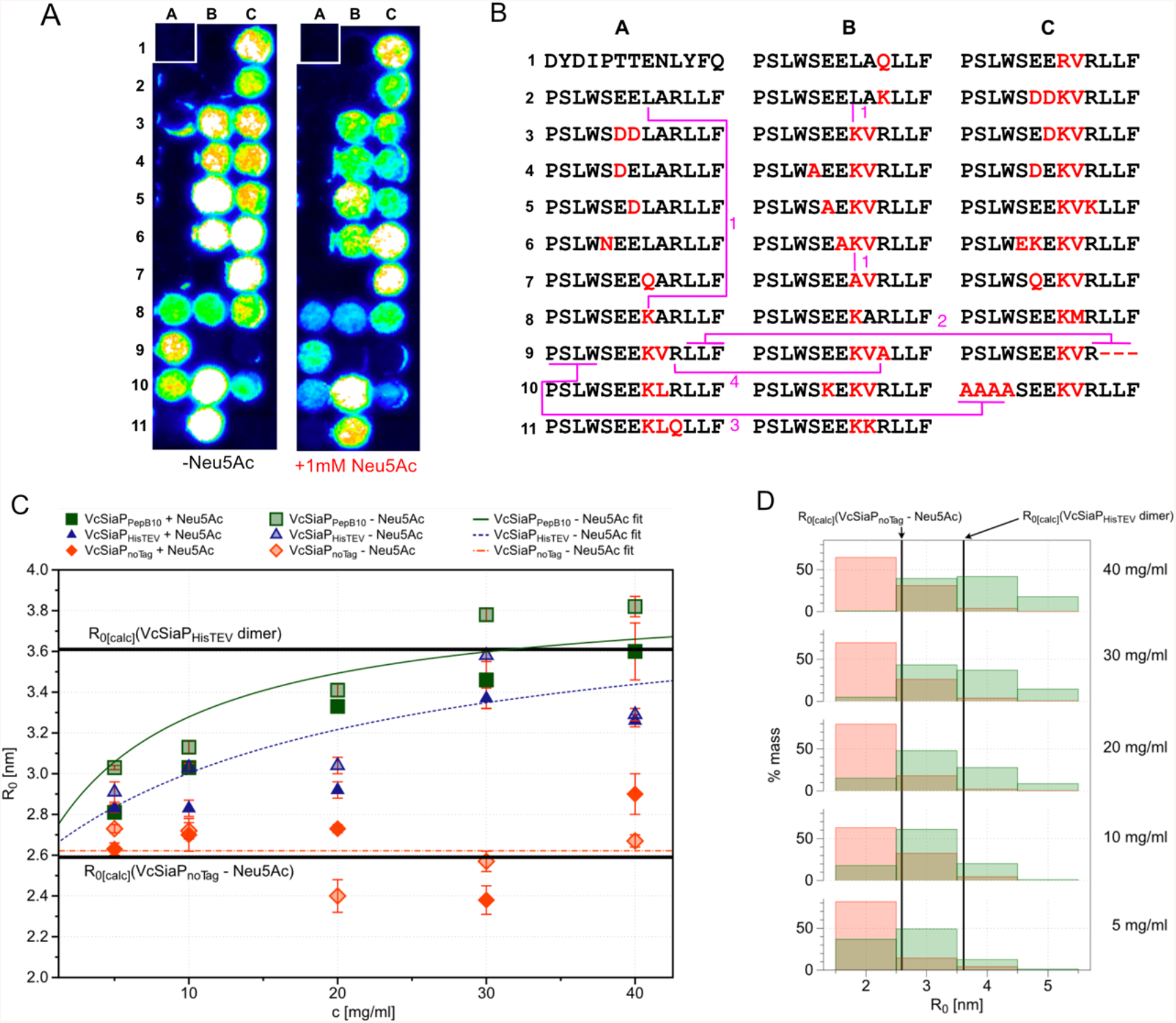
Characterization and strengthening of peptide-VcSiaP interactions. **A)** Spots with synthesized peptides on cellulose membranes after incubation with fluorescence labelled VcSiaP and detection of fluorescence at 800 nm. The experiment was performed with two identical peptide screens, one without addition of sialic acid (left) and one after incubation of VcSiaP with sialic acid (right). **B)** Sequences of the peptide screens from A). Single mutations are indicating in red letters, mutations with clearly different binding of VcSiaP are highlighted in magenta and explained in the main text. **C)** DLS measurements for hydrodynamic radius determination of different VcSiaP constructs at certain concentration. For each protein construct, the measurements were carried out with (dark icons) and without (bright icons) substrate. **D)** Histograms of the radius distributions in mass percent for different concentrations from the untagged protein (red) and the protein with strong binding peptide B10 (green). The calculated radii from the crystal structures are shown as black bars.

Since the tagged SBP is a monomer in gel filtration experiments, we knew that the interaction seen in the crystal lattice is rather weak and the peptide sequence needed to be optimized to see the interaction in solution. Indeed, no fluorescence signal was observed at the position of the tag peptide on the cellulose membrane (Figure 5A, peptide A1). Similarly, negative results were obtained for variations of the tag peptide sequence (not shown). However, inspired by the “scoop-loop” theory for ABC transporters ^30^, we tentatively modeled various periplasmic loops of the QM-domains into the substrate binding cleft by hand and found that the loop connecting TMs 1 and 2 of the Q-domains provided a surprisingly good fit to the binding pocket (SI Figure 4). While this peptide did not bind to the SBP in our experiment (Figure 5A, peptide A2), several variations that we had included in the screen did, resulting in a striking increase in binding strength. For example, simply introducing a positive charge C-terminal of the central double glutamate led to a drastic increase in affinity (peptides A7 vs A8, peptides B2 vs B3 and peptides B6 vs B7 vs B8, magenta 1 in Figure 5AB). In terms of structure, this makes sense, because the introduced lysine residue can potentially interact with E16 deep in the substrate binding pocket (SI Figure 4). The binding could be further strengthened by introducing a second positive charge (peptides B10 and B11 in Figure 6) that can potentially interact with E66 in the SBP. The hydrophobic C-terminal LLF sequence is also important for binding (peptide A9 vs peptide C9, magenta 2 in Figure 5AB). In the crystal structure, this part of the peptide is in contact with a hydrophobic patch on the surface of the SBP. The N-terminal sequence PSLW (peptide A9 vs C10, magenta 3) and the arginine at position 10 of the peptide are also important (peptides B9 vs A9, magenta 4). The latter two features can however not be rationalized by the structural model of the peptide, indicating that the conformation of the peptide in the SBP might change depending on the sequence. To get a rough impression about the strength of the interaction, we performed a competition experiment with 1 mM Neu5Ac using an identical peptide membrane. However, both experiments showed similar or slightly lower fluorescence intensities. Thus, Neu5Ac had an influence on the interaction but due to the very high local peptide concentrations on the membrane, it is difficult to estimate a K_D_ value from such experiments.

Therefore, we further substantiated the peptide binding in solution with dynamic light scattering (DLS) experiments. For this purpose, the sequence connecting the His_6_-tag of VcSiaP was replaced with the sequence of the strong binding peptide identified in the peptide-scan (peptide B10, Figure 5AB). We then performed a series of DLS experiments with different concentrations of both types of tagged (HisTEV and peptide B10) and untagged VcSiaP protein (Figure 5C) and in competition with the natural substrate sialic acid. For the untagged protein, a nearly constant hydrodynamic radius was measured for all different concentrations and, as expected, the addition of sialic acid did not significantly change the observed hydrodynamic radius. Overall, the radii for the untagged samples corresponded reasonably well to the monomeric radius calculated from the crystal structure of the open VcSiaP conformation (Figure 5C). In contrast, for both tagged proteins, we observed a clear dependence of the hydrodynamic radii on the protein concentration (Figure 5CD). At low concentrations, the radii were close to that of the untagged protein, while for higher concentrations the radii increased significantly and converged close to the expected value for the dimeric crystal structure. In line with the peptide-scan experiment, the “optimized tag” led to a stronger increase than the original tag. For both constructs the hydrodynamic radii slightly decreased after addition of sialic acid, maybe because other interactions stabilize the dimer, once it is formed (Figure 4A). Fitting the average hydrodynamic radii of the different concentrations with a binding isotherm allowed us to estimate a K_D_ of ~30 µM for the B10-peptide interaction, which is two magnitudes lower than the affinity towards the sialic acid. It should be noted that the K_D_ for the isolated peptide is likely lower, because the dimerization may entail a cooperative effect.

## Discussion

The conformational dynamics between the open- and closed-states of SBPs have only recently been studied for ABC-transporter related SBPs ^14, 16, 31^ and were completely unknown for TRAP transporter SBPs. Our above described FRET experiments and kinetic analysis reveal that VcSiaP has a ligand dependent substrate-binding rate. Contrary to some SBPs of ABC importers ^15, 17, 32, 33^, no ligand-free closed or ligand-bound open state of VcSiaP were detected in our experiments. Thus, VcSiaP likely uses an induced-fit binding mechanism ^34^. As for ABC importers ^35^, this ligand binding mechanism presumably allows the transmembrane domains of the transporter to discriminate between the ligand-free and ligand-bound SBP and thus to initiate transport, while avoiding futile transport cycles. It should be noted that the time-resolution of ~5 ms used in the present study is not sufficient to fully exclude a mechanism involving very fast intrinsic conformational dynamics < 1 ms or transitions that are so rare that they require a longer observation time to be detected.

The high-resolution crystal structure of the closed-state of VcSiaP presented above completes the basic structural information for the two best studied model systems of TRAP transporter SBPs, VcSiaP and HiSiaP. The tag peptide bound to sialic acid binding site is structurally completely different from the natural substrate. Still, it appears to trigger the same closing mechanism, as evident from the difference distance matrices in Figure 4C. One might therefore speculate, that an important prerequisite for triggering the closing mechanism is a substrate that can physically bridge the gap between the two lobes of the SBP and thereby provides the trigger needed to overcome the open-locked state energetic barrier of SBPs. It is interesting that D_−14_ of the peptide emulates the interaction with natural substrates (Figure 4B). One possible mechanism is therefore that the substrate first binds to a residue that acts as a substrate filter ^21^ and if it is large enough to bridge the gap to the second lobe and make sufficient interactions, the P-domain closes. Small molecules would hence not be able to induce the closed conformation, as exemplified by a sulfate molecule tightly bound to arginine 145 in the open structure of VcSiaP (PDB-ID: 4MAG) ^11^. This fits nicely to the very weak evolutionary coupling between the residues lining the inner surfaces of the gap (Figure 3E). Overall, the structure of the peptide bound VcSiaP protein is reminiscent of the complex between the maltose transporter MalFGK2 and its SBP, were a non-conserved periplasmic loop protrudes into the binding cleft of the SBP ^30^. A similar observation has been made in the complex of the vitamin B12 transporter (BtuCDF) and its SBP ^36^. The feature has been interpreted as a “scoop loop”, which helps to dislocate the bound maltose molecule from the SBP (i.e. the maltose binding protein), or to prevent reassociation of the released ligand with the SBP before it can form a productive interaction with the membrane domains ^30^. It will be interesting to investigate in further studies, if the ability of VcSiaP to bind peptides in the binding site regions are an indication of such a “scoop loop” mechanism for TRAP transporters.

As mentioned above, TRAP transporters are present in many important pathogens such as *V. cholerae* and *H. influenzae*. Since TRAPs are absent in humans, they are potential targets for the development of new antibiotics. The mere fact that a TRAP transporter SBP is able to bind a peptide sequence opens an interesting avenue at developing inhibitors for TRAP transporters, similar to the successful inhibition of vitamin B12 transporter SBP BtuF with a nanobody, bound to the binding site and blocking the substrate binding ^37^. Our peptide screens have shown that with only 32 rationally designed peptides, a strong increase in affinity could be accomplished. With an estimated K_D_ in the range of 30 µM, the tested peptide sequences are still far away from affinities of compounds required for drug development. But, due to the modular structure of peptides, even short sequences give raise to an easily accessible and very large pool of possible structures ^38^. The X-ray structure of the peptide linked SBP dimer immediately suggests that screening approaches such as Bac2Hybrid or phage display assays ^39^ might be efficient ways to find and characterize such inhibitors in further experiments.

## Conclusion

We have used smFRET to analyse the closing mechanism of VcSiaP in real time and found that the closing movement of the TRAP transporter SBP is substrate induced and closely related to SBPs from ABC transporters. The comparison of two ligand-bound structures of VcSiaP, one with the natural and one with a peptide ligand suggests that the closing mechanism of VcSiaP is triggered by physically bridging the gap between the two lobes of the protein. The peptide-bound structure of VcSiaP allowed us to design stronger binding peptides that can serve as starting points to find possible peptide-based inhibitors for TRAP transporters.

## Material & Methods

### Expression and Purification

The proteins were expressed and purified as described before ^40^. The genes for substrate binding proteins VcSiaP and HiSiaP were cloned into a pBADHisTEV vector with a N-terminal 6xHis-TEV tag (Huanting Liu, University of St Andrews). All mutations were performed according to a protocol by Liu et al ^41^. The mutants were expressed in M9-minimal media to prevent the presence of the substrate sialic acid during expression from LB media ^40^. To purify the protein, a Ni^2+^-affinity chromatography was followed by ion-exchange chromatography and size-exclusion chromatography. If necessary, the His-tag was cleaved of by an overnight incubation at 4 °C with 1:50 ratio protein to TEV protease and separated by a second Ni^2+^-affinity chromatography. The purity of protein containing fractions were checked with SDS-PAGE after each step. The purified proteins were concentrated, flash frozen and stored at −80 °C.

### Crystallization and Structure Determination

For crystallization, the purified proteins were concentrated to ~ 25 mg/mL. The crystallization was carried out with vapour diffusion method by using sitting drop 96 well crystallization plates. The substrate bound approaches were previously incubated for 30 minutes on ice with a 2 times excess of sialic acid. In every drop, 0.5 μL protein solution and 0.5 μL screen solution were mixed and the crystal plates were grown for several weeks at room temperature under exclusion of light and vibrations. The drops were regularly checked with a microscope or a protein crystallization imager (Rock Imager 1000, Formulatrix US). For crystallization approaches with substrate Neu5Ac, one crystal was obtained in commercially available screen Morpheus (Molecular Dimensions, UK) well F4 and for the peptide-bound approaches in self-designed optimization screen, based on JCSG-plus (Molecular Dimensions, UK) well A7/D7 (optimized condition: 0.2 M lithium sulfate, 0.1 M Tris (pH 8.5) and 1.26 M ammonium sulfate). The obtained crystals were harvested with a cryo-loop from the initial drop, soak quickly into a mixture of 35 % glycerol reservoir solution for cryo protectant and flash frozen into liquid nitrogen. The diffraction data for VcSiaP dimer were collected at beamline BL14.3 of BESSYII (Berlin, Germany), with a MarMOSAIC 225 CCD detector. The data for VcSiaP with sialic acid were collected at beamline PX1 (Zuerich, Switzerland) with EIGER 16M detector. The diffraction data were integrated with XDS ^42^ and the structures were solved by PHASER ^24^ with molecular replacement using PDB-ID 4MAG ^11^ for VcSiaP dimer and PDB-ID 3B50 for sialic acid bound VcSiaP ^10^. The structures were refined with PHENIX ^43^ and COOT ^44^ and the quality of the model was checked after each step with MolProbity ^45^.

### ITC measurement

The ITC experiments were performed with a MicroCal PEAQ-ITC machine from Malvern Panalytical (UK). Previously to every measurement, the sample cell was equilibrated three times with 300 μL standard protein buffer (50 mM Tris, 50 mM NaCl, pH=8). The protein solution was diluted to 100 μM in a total volume of 350 μL with standard buffer. After transfering the protein solution into the sample cell without any air, the concentration of the remaining solution was determined via UV-Vis absorption with a NanoDrop 2000 (ThermoFisher Scientific, US). The syringe was automatically loaded with 1.2 mM sialic acid in standard protein buffer. The experiment was carried out isothermaly at 25 °C and 19 injections of the ligand with a volume of 2 μL per injections were titrated into the sample cell. The experimental data were recorded and analysed with commercial software from Malvern Panalytical. The heat pulses were integrated and the isotherm curve for each measurement were fitted with the one set of sites fitting mode to get the experimental and thermodynamic parameters.

### Protein labelling

Stochastic labelling with the dyes Alexa555 and Alexa647 maleimide (ThermoFisher) was performed as done previously ^16^. In brief, the purified proteins were first treated with 10 mM Dithiothreitol (DTT) for 30 min to fully reduce oxidized cysteines. After dilution of the protein sample to a DTT concentration of 1 mM the reduced protein was immobilized on a Ni^2+^-Sepharose resin and washed with ten column volumes of buffer A (50 mM Tris-HCl, pH 7.4, 50 mM KCl). The resin was incubated 1-8 h at 4 °C with a dye-to-protein ratio of ~20 dissolved in buffer A. Subsequently, unbound dyes were removed by washing with twenty column volumes of buffer A. Elution of the proteins was done with buffer A and 400 mM imidazole. The labelled proteins were further purified by size-exclusion chromatography (Superdex 200, GE Healthcare) using buffer A.

### Solution-based smFRET experiments

Solution based smFRET experiments ^46^ were performed on a homebuilt confocal ALEX ^47^ microscope as previously described ^16, 17^. All sample solutions were measured with 100 μl drop on a coverslip with concentration of around 50 pM in buffer A.

The fluorescent donor molecules were excited by a diode laser OBIS 532-100-LS (Coherent, USA) at 532 nm operated at 60 μW. The fluorescent acceptor molecules are excited by a diode laser OBIS 640-100-LX (Coherent, USA) at 640 nm operated at 25 μW at the sample in alternation mode (50 μs alternation period). The lasers were combined by a polarization maintaining single-mode fiber P3-488PM-FC-2 (Thorlabs, USA). The laser light is guided into the epi-illuminated confocal microscope (Olympus IX71, Hamburg, Germany) by dual-edge beamsplitter ZT532/640rpc (Chroma/AHF) focussed by a water immersion objective (UPlanSApo 60x/1.2w, Olympus Hamburg, Germany). The emitted fluorescence is collected through the objective and spatially filtered using a pinhole with 50 μm diameter and spectrally split into donor and acceptor channel by a single-edge dichroic mirror H643 LPXR (AHF). Fluorescence emission was filtered (donor: BrightLine HC 582/75 (Semrock/AHF), acceptor: Longpass 647 LP Edge Basic (Semroch/AHF) and focused on avalanche photodiodes (SPCM-AQRH-64, Excelitas). The detector outputs were recorded by a NI-Card PCI-6602 (National Instruments, USA).

Data analysis was performed using home written software package as described in ^14^. Single-molecule events were identified using an All-Photon-Burst-Search algorithm with a threshold of 15, a time window of 500 μs and a minimum total photon number of 50 ^48^. Photon count data were extracted, background subtracted and plotted in ES-histogram with a minimum of 150 counts.

E-histogram of double-labelled FRET species with Alexa555 and Alexa647 was extracted by selecting 0.25<S<0.75. E-histograms of open state without ligand (apo) and closed state with saturation of Neu5Ac at 200 μM concentration (holo) were fitted with a Gaussian distribution 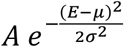 resulting in (*μ*_0_ = 0.866; *σ*_0_ = 0.059) and (*μ*_c_ = 0.932; *σ*_c_ = 0.031) for open and closed state. The closed fraction in the titration experiment was calculated as 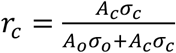 from the double Gaussian fit to the data 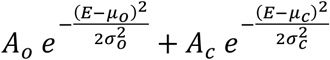, where mean and standard deviations were fixed to the respective stated values.

### Surface-based smFRET experiments

Confocal scanning experiments were performed at room temperature and using a home-built confocal scanning microscope as described previously ^14^. In brief, surface scanning was performed using a XYZ-piezo stage with 100 Ч 100 Ч 20 μm range (P-517–3 CD with E-725.3CDA, Physik Instrumente). The detector signal was registered using a HydraHarp 400 picosecond event timer and a module for time-correlated single photon counting (both Picoquant).

Surface immobilization was done as follows. Microscope cover-slides (No. 1.5, Marienfeld) were cleaned by sonication in ethanol, acetone and MQ water for 10 min each, followed by plasma etching (Plasma Etch, PE-25-JW) for 10 min. Functionalization with PEG-Silane (6-9 PE units) and Biotin-PEG-Silane (MW3400, Laysan Inc.) was done in toluene (55 °C; o/n). The surfaces were incubated for 10 min with 0.2 mg/ml neutravidin (Invitrogen) in buffer A (50 mM Tris-HCl, pH 7.4, 50 mM KCl). Unbound neutravidin was washed away with buffer A. Next, the surface was incubated with 1 nM biotinylated penta-His antibody (Qiagen) in buffer A for 10 min. Then, labelled proteins were immobilized by incubating with 5-100 pM of labelled protein in buffer A for 5 min. Unbound proteins were washed away with buffer A.

All surface-based smFRET measurements were done as described previously ^16, 17^. In brief, measurements were carried out in buffer A supplemented with 1 mM of 6-Hydroxy-2,5,7,8-tetramethylchromane-2-carboxylic acid (Trolox) and 10 mM Cysteamine. Data were recorded with constant 532 nm excitation at an intensity of 0.5 μW (~125 W/cm^2^). Scanning images of 10 × 10 μm were recorded with 50 nm step size and 2 ms integration time at each pixel. After each surface scan, the positions of labelled proteins were identified manually. The position information was used to subsequently generate fluorescence trajectories with a time resolution of 5 ms. The donor and acceptor fluorescence counts were used to calculate the apparent FRET efficiency, via F_DA_/(F_DA_+F_DD_), were F_DA_ and F_DD_ are the uncorrected acceptor and donor photon counts. The apparent FRET efficiency time-traces were further analysed with hidden Markov modelling as described previously ^17^. In the absence of ligand and in the presence of 1 mM Neu5Ac the time-traces were fitted with one state, and in the presence of 500 nM Neu5Ac with two states. From the fit, the individual lifetimes could be extracted as well as the average apparent FRET efficiency of each state.

### Peptide Screens

To detect the interaction of VcSiaP with several peptides on the screen surface, a construct VcSiaP Q245C was expressed and purified as described before. To avoid dimerization of the introduced cysteine, every buffer was supplemented with 1 mM TCEP (Tris(2-carboxyethyl)phosphine). Before labelling, the TCEP was removed through a size-exclusion chromatography step on a SD 200 10/300 with standard protein buffer without TCEP. Immediately after elution, the protein fractions were combined and supplemented with four times excess of DyLight800 Maleimide Fluorophore Label (ThermoFisher Scientific, US). The solution was incubated overnight under light exclusion at 4 °C. On the next day, the remaining free label was removed with a PD-10 column and standard protein buffer. The protein elution fraction was concentrated up to 1 mg/mL.

The peptide screen was designed based on the interaction in the VcSiaP dimer structure and ordered from peptides&elephants (Germany). The peptides were synthesized in single spots on a cellulose membrane. Before incubation with the target protein, the membrane was soaked with water, washed three times for 15 seconds with methanol and again soaked into water. For reduction of unspecific binding, the peptide screen was incubated for 1 h at room temperature with 1 % BSA in standard protein buffer (50 mM Tris, 50 mM NaCl, pH=8). After three washing steps with buffer A for each 10 minutes the screen was incubated for 1 h at room temperature in 10 mL buffer A with 1 μg VcSiaP Q245-DyLight800. Afterwards the screen was again washed three times for 10 minutes with buffer A. The fluorescence onto the screen was detected with an Odyssey Imaging System at a wavelength of 800 nm.

### DLS measurements

For the determination of the hydrodynamic radius R0 of different VcSiaP constructs, protein solutions of different concentrations were prepared for each construct in the same manner. The purified protein solution was concentrated to a final concentration of 40 mg/ml. Subsequently, to one half of the volume 100 mM sialic acid solution (dissolved in 50 mM NaCl, 50 mM Tris, pH=8) was added to a final concentration of 10 mM. The other half of the concentrated protein solution was diluted with 1/10 of buffer (50 mM NaCl, 50 mM Tris, pH=8). Both of the solutions were then diluted step-wise to result in a concentration series with protein concentrations of 40 mg/ml, 30 mg/ml, 20 mg/ml, 10 mg/ml, and 5 mg/ml. These solutions were centrifugated at 15000 x *g* for 15 min to remove aggregates and measured in a DLS-device (DynaPro® NanoStar® (Wyatt Technology Corporation)). For each condition, three measurements of 10 μl of VcSiaP solution were done at a sample temperature of T = 25 °C and three measurement cycles of each 20 single data acquisitions with acquisition times of T_acq_ = 3 s. For the calculation of the hydrodynamic radii based on crystal structures, the online tool ‘HullRad’ ^49^ was used (R(dim)=3.61 nm, R(monomer, apo)=2.59 nm, R(monomer, holo)=2.53 nm).

## Accession numbers

PDB: 7A5Q

PDB: 7A5C

## Abbreviations

SBP: Substrate binding protein
TRAP: tripartite ATP-independent periplasmic

## Glossary

TRAP transporters: Tripartite ATP-independent periplasmic transporters
SBP: Substrate binding protein
ABC transporter: ATP binding cassette transporter

## Acknowledgements

This project was financed by the German Research Foundation (Deutsche Forschungsgemeinschaft, DFG) in project no. HA 6805/5-1 (to GH), GRK2062 – project number C03 – (to T.C.), SFB 863 – Project number A13 – 111166240 (to TC) and the European Research Council (ERC-StG “SM-IMPORT”, No. 638536 to TC). MFP acknowledges a PhD fellowship from the Konrad Adenauer Stiftung. CG acknowledges a PhD fellowship from the Studienstiftung des deutschen Volkes. GH thanks Olav Schiemann (University of Bonn) for support. GH thanks Matthias Geyer (University of Bonn) for helpful discussions. JG and GH thank Anton Schmitz (caesar, Bonn) for help with the fluorescence detection of the peptide screens. The authors thank beamline staff at SLS (Zuerich) and BESSY (Berlin) for support during data collection.

## Author contributions

GH conceived this study. MFP, CG, JG, GHT, TC and GH have designed the experiments and analyzed the data. MFP and NS expressed and purified the protein constructs and determined the protein structures, JG performed the peptide scan experiments. CG and MdB performed single-molecule FRET experiments. MFP and GH wrote the manuscript with input from all authors.

## Supporting information

**SI Figure 1:**
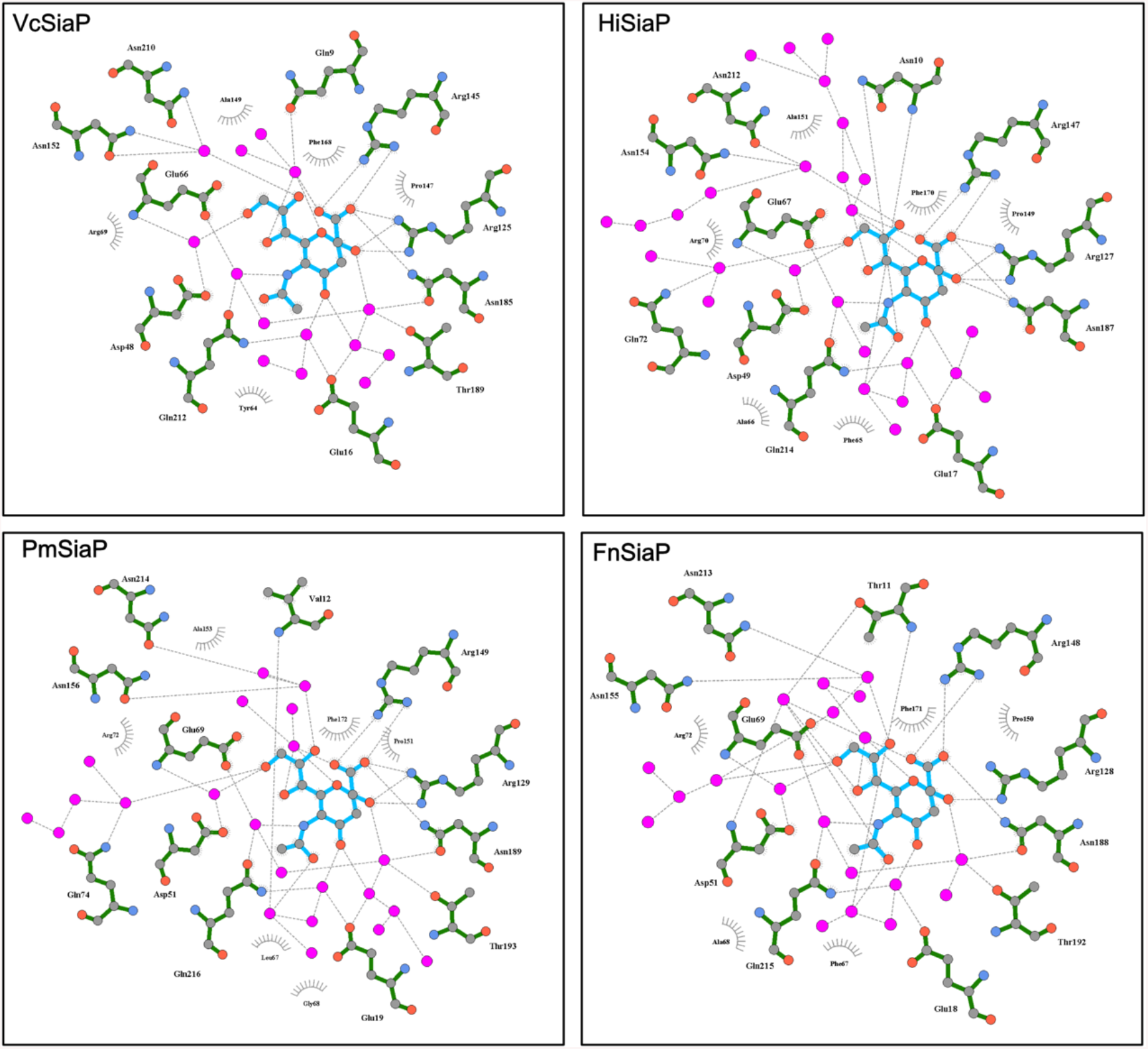
Interaction between sialic acid and TRAP SBPs. The substrate interaction interfaces were calculated with *LigPlot* ^50^ and the corresponding sialic acid-bound SBPs (VcSiaP from *Vibrio cholerae* (this work), HiSiaP from *Haemophilus influenzae* (PDB-ID: 3B50 ^10^), PmSiaP from *Pasturella multocida* (PDB-ID: 4MMP ^11^) and FnSiaP from *Fusobacterium nucleatum* (PDB-ID: 4MNP ^11^)

**SI Figure 2:**
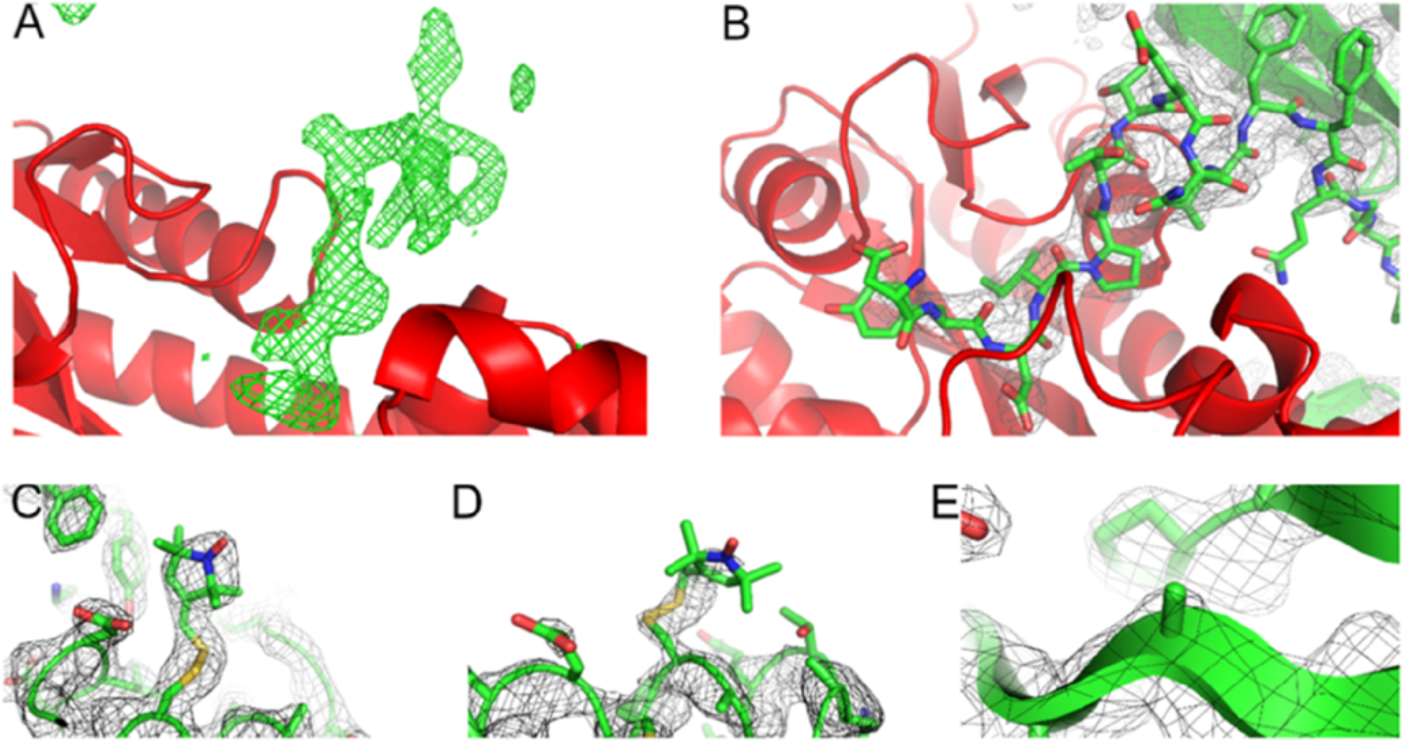
Structure refinement and comparison of peptide-VcSiaP interaction. **A)** Difference electron density (green, σ=5.0) reveals a clear peptide-chain like discrepancy in the active binding of VcSiaP (cartoon, red). **B)** The N-terminal affinity tag could be successfully built into the difference electron density from A) and connect to a neighboured VcSiaP monomer (σ=1.5). **C)** The spin label MTSSL attached to a cysteine at position Q54C (σ=1.5). **D)** The spin label MTSSL attached to a cysteine at position L173C (σ=1.5). **E)** Mutation R125A from previous studies to validate the overall protein structure for this mutant.

**SI Figure 3:**
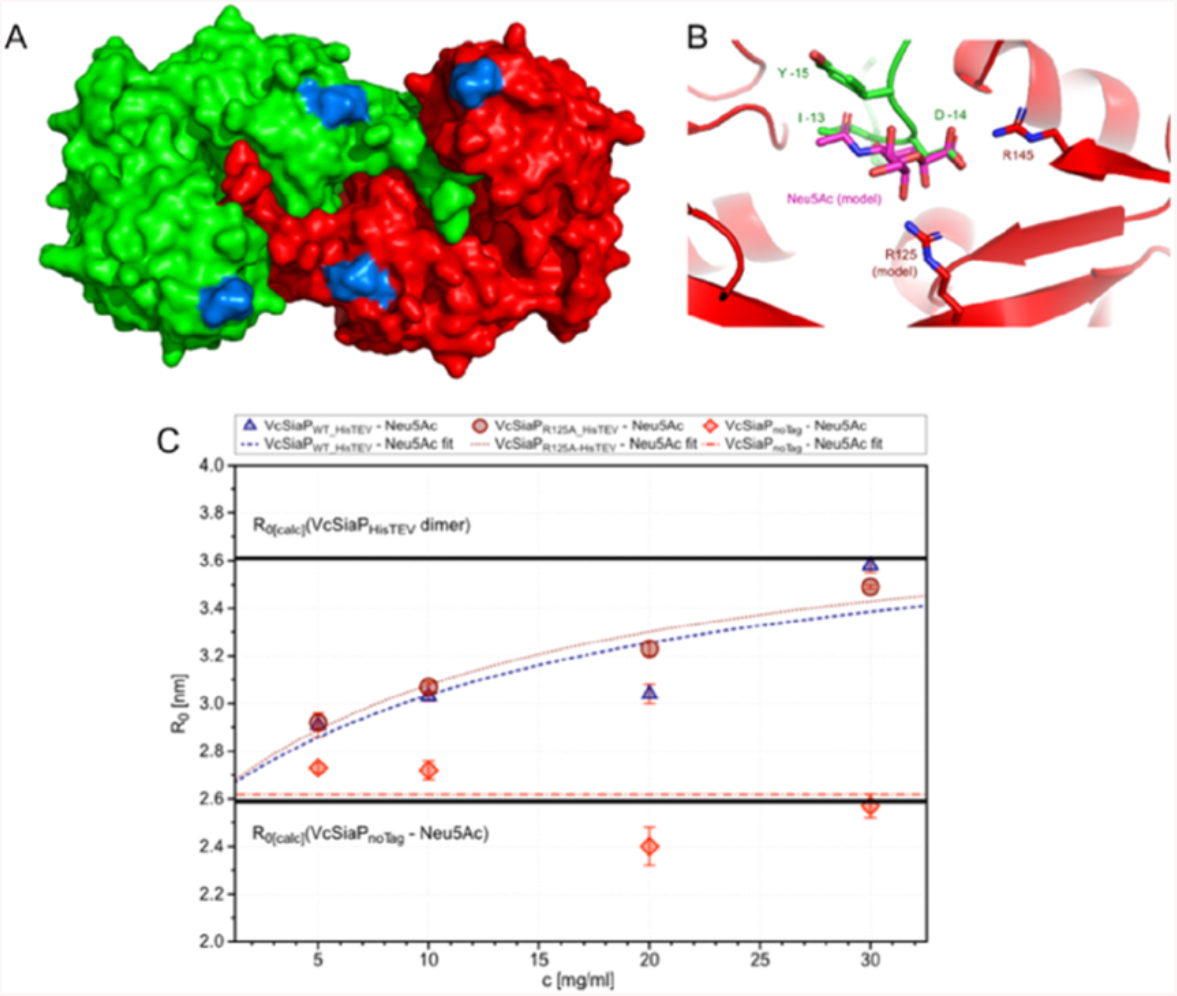
Analysis of the dimeric VcSiaP interface. **A)** Surface representation of the dimeric VcSiaP with affinity tag bound to the active binding site of the opposite VcSiaP. One chain is represented in green, the other one in red. The spin label positions at positions Q54C and L173C are represented in blue. **B)** Modelling of Arg125 back into the crystal structure of the dimer showing no obvious interruptions into peptide binding. For a comparison of artificial peptide and natural substrate, the sialic acid (magenta) was modelled into the dimeric protein structure. **C)** Hydrodynamic radii from DLS experiments for VcSiaP wildtype with HisTEV (dark red), VcSiaP wildtype without HisTEV (light red) and VcSiaP R125A Q54R1 L173R1 with HisTEV (blue), which was used for crystallization of VcSiaP dimer.

**SI Figure 4:**
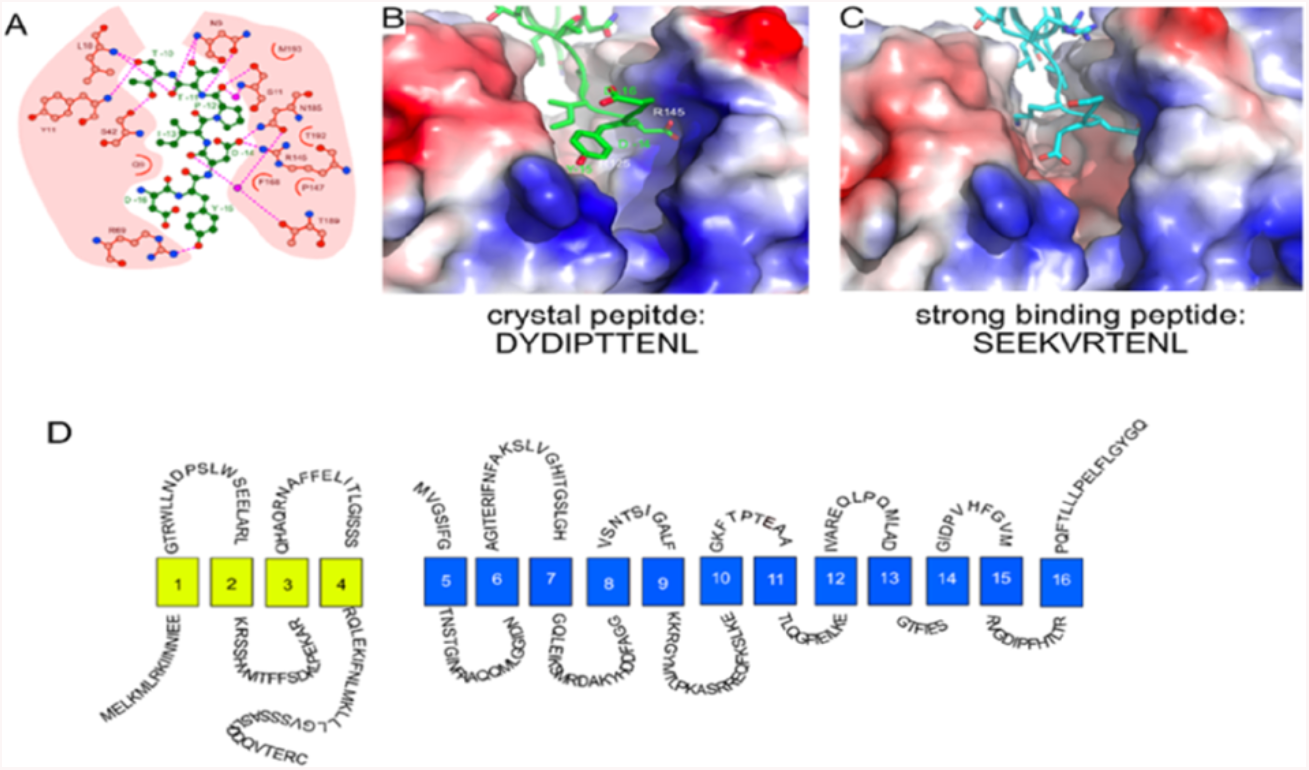
Characterisation of the interaction between peptides and VcSiaP active binding site. **A)** Analysis of the peptide-active binding site interaction with LigPlot ^50^. The peptide is shown in green, the amino acids of the active binding site in pink. Strong interactions are represented as dashed, pink lines. **B)** Representation of the peptide inside the active binding cleft of VcSiaP. The peptide is shown in green sticks, the surface of VcSiaP with electrostatics. **C)** Same as B) but with model of a strong binding peptide (cyan sticks) from Figure 5 A9. **D)** Hypothetical profile of the membrane domains Q (yellow) and M (blue) from the TRAP transporter VcSiaPQM, the C-terminus be located on the cytoplasm side and the N-terminus on the periplasm site ^7^.

## Notes

### Competing Interest Statement

The authors have declared no competing interest.

